# Divergent 3D genome organization in livers of cave and surface morphs of *Astyanax mexicanus* as a potential driver of unique metabolic adaptations in cave environment

**DOI:** 10.1101/2024.09.30.615929

**Authors:** Tathagata Biswas, Hua Li, Nicolas Rohner

## Abstract

The cave morphs of *Astyanax mexicanus* have evolved a suite of distinct adaptations to life in perpetual darkness, including the loss of eyes and pigmentation loss, as well as profound metabolic changes such as hyperphagia and starvation resilience, traits that sharply contrast with those of their river-dwelling surface counterparts. While changed gene expression is a primary driver of these adaptations, the underlying role of 3D genome organization – a key regulator of gene expression – remains unexplored. Here, we investigate the 3D genome architecture of the livers of surface fish and two cavefish morphs (Pachón and Tinaja) using Hi-C, performing the first comparative 3D genomic analysis in this species. We analyzed and identified cave-specific 3D genomic features, such as genomic compartments and loops, which were conserved in both the cave populations but absent in surface fish. Integrating the 3D genome data with transcriptomic and epigenetic datasets, linked these changes to differential expression of metabolically relevant genes, such as *Arhgef19* and *Endog*. Additionally, our study also uncovered genomic inversions unique to cavefish, potentially tied to cave adaptation. Our findings suggest that 3D genome organization contributes to transcriptomic shifts underlying cavefish phenotypes, providing a novel intra-species and morph specific perspective on 3D chromatin evolution. This study establishes a foundation for exploring how genome architecture potentially facilitates adaptation to new environments. Comparison of morphs within the same species also establishes a foundation for better understanding of how 3D genome reorganization may drive speciation and phenotypic diversity.

## Introduction

The fish *Astyanax mexicanus* exists in two distinct morphs: river-dwelling surface fish and multiple parallely evolved cave-dwelling cavefish ^1, 2^. Cavefish, having adapted to the dark and nutrient-scarce cave environment, exhibit striking traits such as eye and pigment loss, along with metabolic adaptations such as resilience to starvation, enhanced lipogenesis, persistent hyperglycemia, and hyperphagia ^3-9^. Comparative molecular studies have shown that these metabolic differences between surface and cavefish are largely driven by gene expression changes^10, 11^. While prior research has explored the roles of both cis- and trans-regulatory elements in shaping metabolic and transcriptomic changes in *A. mexicanus*, the role of three-dimensional (3D) genome organization – a critical regulator of gene expression – has remained unexplored.

Within the nucleus, 3D genome organization governs the transcriptomic profile by determining the spatial proximity of regulatory elements, such as enhancers, to their target genes ^12-14^. This organization involves genome looping, where DNA segments spanning a few to hundreds of kilobases fold to bring distant regulatory elements into contact with genes, a process facilitated by cohesin and CTCF-associated protein complexes through loop extrusion ^15-17^. By allowing genes separated by large genomic distances to share common regulatory elements, these loops play a crucial role in gene regulation ^18, 19^. Beyond these fine-scale structures, the genome is organized into broader active (A) and inactive (B) compartments based on transcriptional activity, gene density, and looping interactions ^20^.

While some studies suggest that gene expression remains relatively robust against changes to the 3D genome organization ^21, 22^, it is widely recognized that alterations in genome architecture can impact molecular machinery, ultimately affecting transcriptional landscapes and even contributing to disease states ^23-25^. Given these effects, understanding the stability of 3D genome organization across evolutionary timescales becomes essential. Research has shown that 3D genome architecture is remarkably conserved over evolutionary timescales, with syntenic regions often maintaining structural integrity alongside consistent gene expression across species ^26-29^. Nevertheless, species-specific variations in genome structure do occur and have been linked to differences in gene expression between species ^27-31^. However, much less is known about how 3D genome architecture may vary within a species, particularly between morphs that have adapted to contrasting environments. If such morph-specific differences exist, they could provide a direct link between variations in genome architecture and adaptive phenotypic divergence.

To explore this, we investigated the 3D genome architecture of *A. mexicanus* surface fish and two cavefish morphs – Pachón and Tinaja – focusing on the liver due to its central metabolic role and the availability of extensive transcriptomic and epigenetic datasets for this tissue (Fig. 1a). Using Hi-C, a technique for mapping genome-wide chromatin interactions, we generated the first 3D genome contact maps for this species. Utilizing the two cavefish morphs, we identified morph-specific chromatin loops, compartmental changes, and shared structural variants affecting genes involved in metabolism and stress response, positioning 3D genome organization as a previously overlooked contributor to metabolic adaptation in response to environmental pressures.

**Figure 1:**
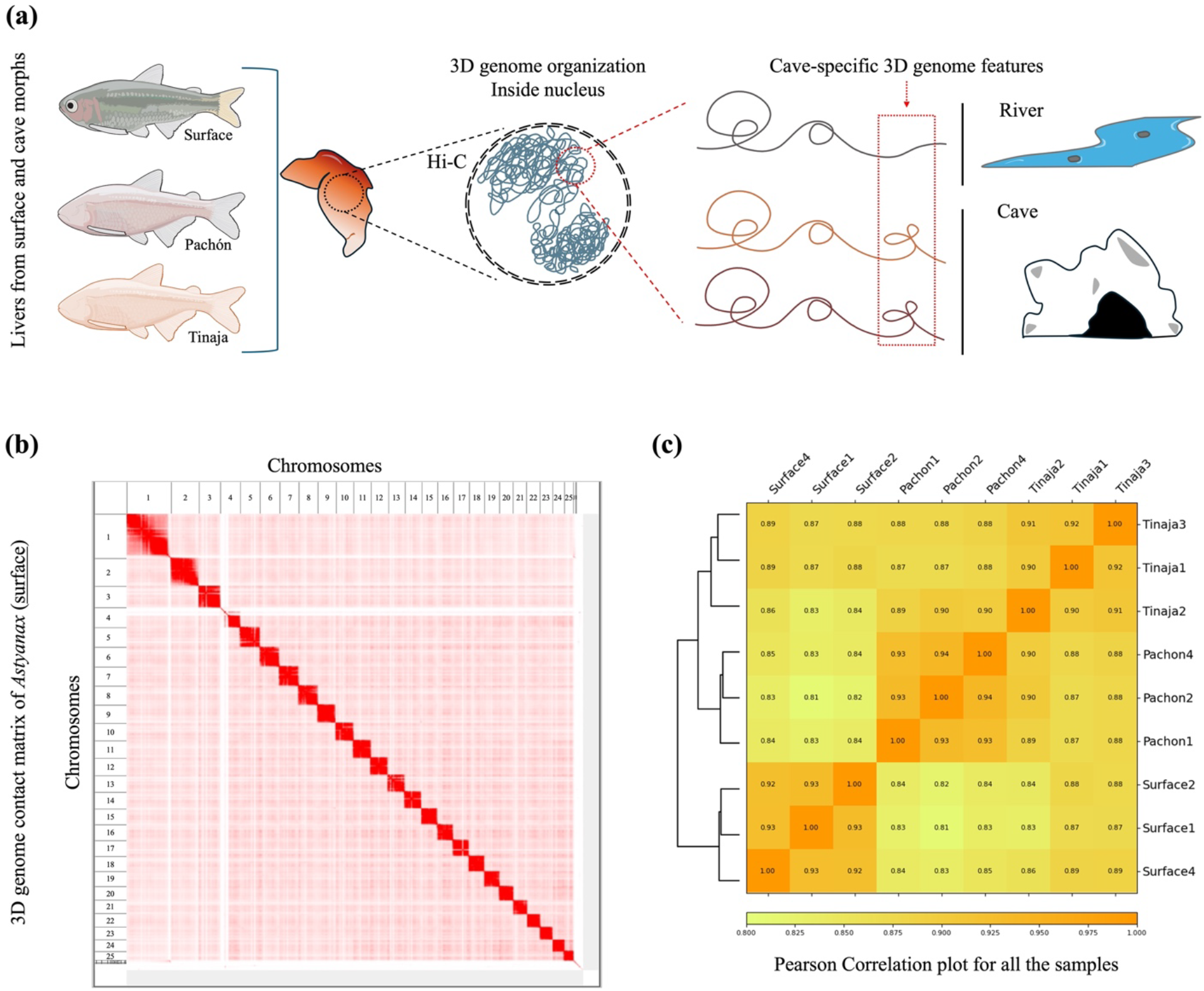
(a) Hi-C experimental plan on livers of surface and cave morphs of *Astyanax mexicanus* to investigate the changes in 3D genome organization within the same species but living under different environmental conditions. (b) Whole genome 3D contact matrix of *A. mexicanus* (surface) liver derived from Hi-C data. (c) Pearson correlation summary for the Hi-C samples sequenced, showing high degree of correlation in-between each of the morphs and biological replicates.

## Materials and Methods

### Sample collection for Hi-C

Liver tissues were collected from three adult fish from each morph: Pachón (499 days, 1M, 2F), Tinaja (511 days, 1M, 2F), and surface fish (553 days, 3F). After a quick 1xPBS wash, whole livers were flash frozen in liquid nitrogen. The frozen tissues were homogenized with help of Zirconium beads (3mm) in a Beadbug 6 microtube homogenizer at 4000 oscillations/minute for 30secs. Thereafter, Hi-C data was generated using the Arima-HiC kit (PN: A510008), which streamlined the protocol as it included reagents optimized for efficient chromatin digestion, proximity ligation, and DNA purification. After necessary QC, sample were ready for library preparation.

### Libraries and Sequencing

The Pachón and surface fish data were generated in the first round. Hi-C libraries were prepared according to manufacturer’s directions for the Arima-HiC Kit: Library Preparation using KAPA Hyper Prep Kit protocol (Document Part No. A160139 v00) starting with 1.95-5.59ug of large proximally-ligated DNA molecules for Pachón and surface, and with 0.56-0.81 ug of large proximally-ligated DNA molecules for Tinaja, as assessed using the Qubit Fluorometer (Life Technologies). Libraries were prepared using the S220 Focused-ultrasonicator (Covaris) to shear samples to 400bp, and the KAPA Hyper Prep Kit (Roche Cat #: KK8500/7962312001) with Illumina IDT-TruSeq DNA UD Indexes (Illumina Cat #: 20020590) following Arima’s modified protocol with 10 cycles of library PCR amplification. Resulting short fragment libraries were checked for quality and quantity using the Bioanalyzer (Agilent) and Qubit Fluorometer (Life Technologies). Libraries were pooled, re-quantified and sequenced as 150bp paired reads on an S1 flow cell (v1.5) on the Illumina NovaSeq 6000 instrument (Surface and Pachón) and P3 flow cell on the Illumina Nextseq 2000 instrument (Tinaja), utilizing RTA and instrument software versions current at the time of processing.

### HiC-seq data process

The Hi-C experiment reads were processed using Juicer from Aiden lab with the default parameters ^32^, against the reference genome AMEX_1.1. Final contact databases were created using mapq >= 30. A total of 620 million to 830 million reads were obtained per sample. Following quality filtering, the number of valid read pairs ranged from 189 million to 419 million.

### A/B compartment analysis

A/B compartment analysis was performed at 25kb resolution using POSSUMM ^33^. The choice of the first or the second PCA (Principal Component Analysis) depended on how it corresponded to the plaid-like pattern observed in the Hi-C data. Once the PCA was decided, the signs of these values were determined to make sure that the positive values always correspond to the active state, which is represented by higher gene density as compared to the inactive state. The significant A/B bin switches were identified using lm() function in base R package. We intersected the gene model with A/B bins using bedtools intersect.

### Loop analysis using SIP method and CTCF motifs

Individual sample data (each .hic file) within each morph were merged to form one dataset (one merged .hic file) for each of the three *Astyanax* morphs. We used the SIP method to detect the genomic loops from the data. Loops were identified using the default parameters with 5k, 10k, 15k, and 20k resolutions. Results were then combined to generate a final loops file for each sample. Loops that are wider than 1MB were excluded. Additionally, MotifFinder was used to find putative CTCF motifs in the anchor points of the identified loops. MotifFinder: https://github.com/aidenlab/juicer/wiki/MotifFinder.

### Aligning other sequencing data with Hi-C

ATAC-seq, Chip-seq and RNA-seq data were downloaded from GSE153052. ATAC-seq data was aligned to AMEX_1.1 using bowtie2. MACS2 and then IDR was used to call peaks for each replicate. Reads were aligned to AMEX_1.1 genome and the gene model was mapped from astMex2 Ensembl106 gene model, using liftoff ^34^ and STAR. The gene expression values TPM were estimated using RSEM. TPMs were normalized using the quantile normalization method to remove any potential batch effect. Differential gene expression analysis was done using R package limma ^35^.

## Results

### Hi-C based 3D genome organization in surface fish, Pachón, Tinaja

We generated Hi-C-based contact matrices for liver tissues of adult surface fish, Pachón, and Tinaja following the Arima-HiC protocol. After sequencing and merging biological replicates, we obtained 1,348 million valid Hi-C reads for surface fish, 1,093 million for Pachón, and 603 million for Tinaja, averaging 304 million contacts per morph. These data enabled high-resolution contact matrices (up to 5 kb), revealing clear delineation of the 25 chromosomes in each morph, with strong intra-chromosomal interactions (Fig. 1b). Notably, being still the same species, the surface and cave morphs exhibited an overall good correlation of the liver Hi-C sequence data (Pearson’s correlation 0.81 ≥ R ≥ 0.89). Also, as expected, biological replicates within each morph of *A. mexicanus* showed a higher degree of correlation (Pearson’s correlation 0.90 ≥ R ≥ 0.94) (Fig. 1c).

### 3D genome segregation into active (A) and inactive (B) compartments

Next, we analyzed the segregation of the genome into A and B compartments, a hallmark of 3D chromatin organization observed across species. Genomic regions with higher gene density and transcriptional activity tend to cluster in the ‘A’ compartment, while less active loci tend to fall within the ‘B’ compartment ^20^. This compartmentalization is visually evident as a plaid-like pattern in Hi-C contact matrices. Using Principal Component Analysis (PCA) Of Sparse, SUper Massive Matrices (POSSUMM) ^33^, we identified A/B compartments in the genome of liver cells of the three morphs of *Astyanax mexicanus* (Fig. 2a, Supplementary Fig. 1). We validated these by integrating liver RNA-seq data ^11^ which confirmed that genes in the A compartment exhibited significantly higher expression levels than those in the B compartment, consistent with their functional distinctions (Fig. 2b).

**Figure 2:**
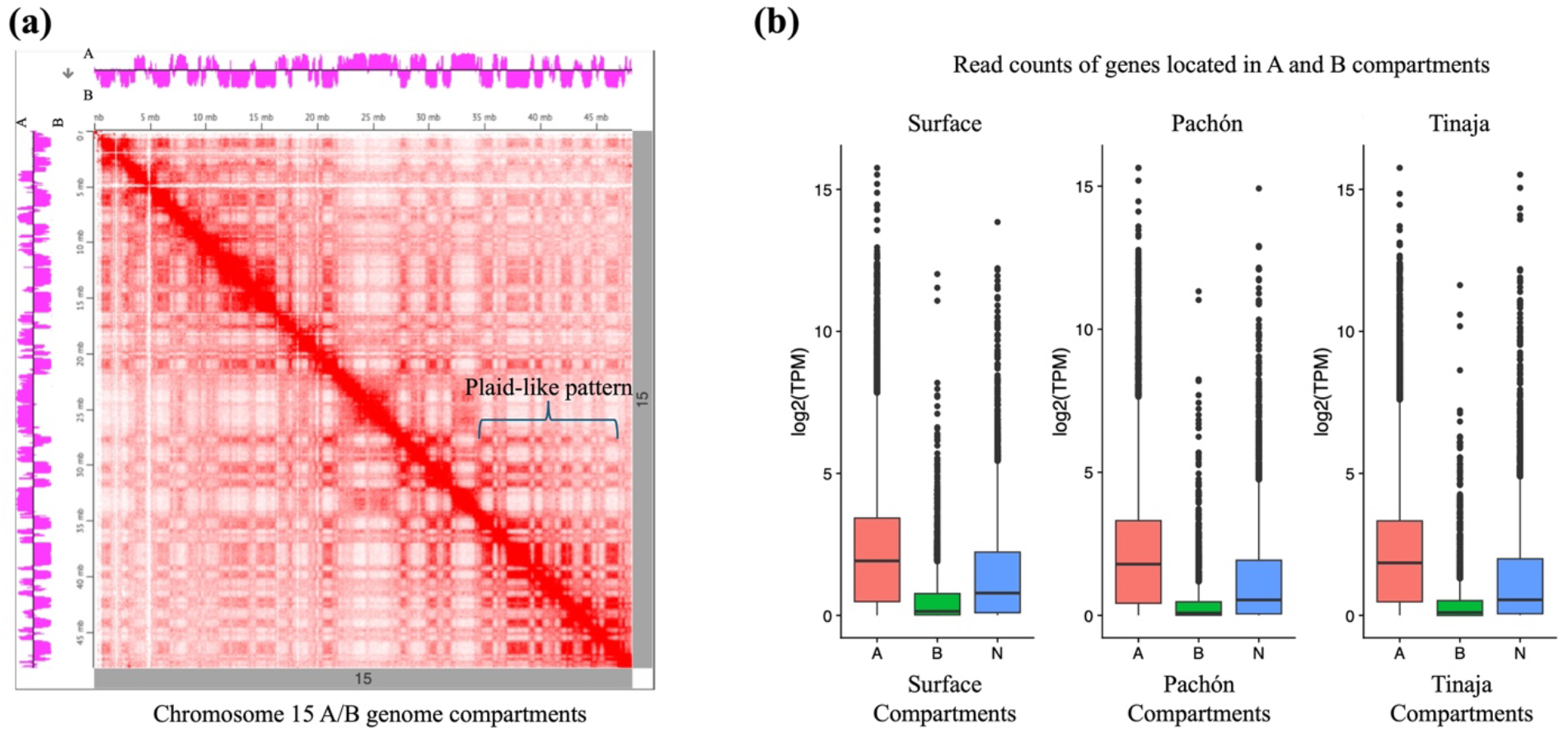
(a) Plaid-like patterns in the chromosomal interaction matrix (chromosome 15) are observed in Pachón, indicating A/B compartmentalization. The magenta track adjacent to the axes represents A/B compartments, aligning with the plaid-like pattern. Positive values correspond to A compartments (active, gene-rich regions), while negative values indicate B compartments (inactive, gene-poor regions). (b) Gene residing in the active ‘A’ compartment show higher transcriptional activity across all the three morphs of *A. mexicanus* as compared to genes residing in the ‘B’ compartment. N denotes the neutral compartment.

### Switching of the genome compartments in cavefish

Given that cave adaptation involves extensive transcriptomic and regulatory remodeling ^6, 10, 11^, we next asked whether changes in compartment status contribute to this divergence. Comparing the morphs, we found that while compartmentalization was largely conserved, 665 out of 53,988 genomic bins (25 kb each) exhibited A/B switching in each of the cavefish samples relative to surface fish (p ≤ 0.001). Genes in regions switching to the A compartment in cavefish showed higher transcriptional fold changes than those switching to the B compartment (Fig. 3a, Supplementary Fig. 2). In contrast, genes located in regions that remained in the same compartment across both cave morphs and surface fish showed no significant transcriptional changes. These findings suggest that large-scale chromatin reorganization supports transcriptomic shifts during cave adaptation.

**Figure 3:**
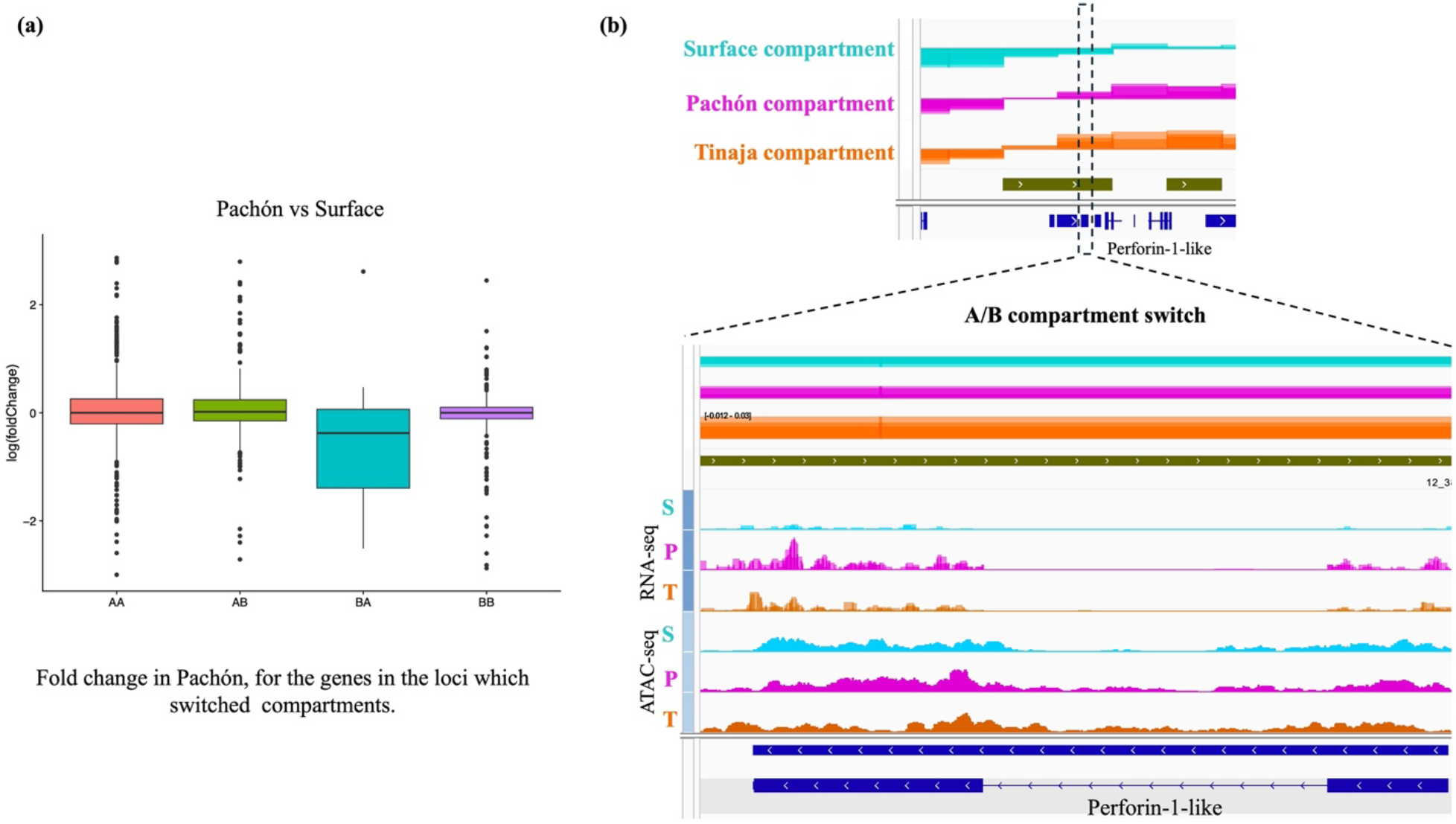
(a) Boxplot depicting fold change in expression for the genes in the loci which switched compartments (Pachón vs surface depicted in the figure). AA and BB denotes unchanged compartments, AB denotes loci which is active in Pachón and inactive in surface, BA denotes loci active in surface and inactive in Pachón. (b) Example of regions that switched from the B compartment in surface fish to the A compartment in both Pachón and Tinaja cavefish (following the order: surface, Pachón, Tinaja; termed BAA regions). The switched BAA compartment loci include genes such as Perforin-1-like (dashed inset), which show higher expression in both cavefish morphs compared to surface fish when aligned with the RNA-seq profile. S is surface, P is Pachón and T is Tinaja.

### Emergence of cave-specific compartmentalization in *Astyanax mexicanus*

If the compartmental reorganization of the genome in the liver contributes to the transcriptomic changes necessary for cave adaptation, we can assume that a set of A/B switches in *Astyanax* livers are critically important for this adaptation. Consequently, to test whether these compartmental switches were linked to cave adaptation, we sought switches common to both Pachón and Tinaja but absent in surface fish – termed cave-specific genome compartmental switches.

Comparing liver genome compartments across the three morphs, we identified multiple compartmental switches that were common between Pachón and Tinaja cavefish when compared to surface fish. We identified regions switching from B in surface fish to A in both cave morphs (BAA regions), containing 287 genes (an example in Fig. 3b), and from A to B (ABB regions), containing 36 genes (Supp. Table 1). To assess whether the cave-specific genome compartment shifts from inactive B compartments (in surface fish) to active A compartments (in cave morphs) – referred to as BAA transitions – are linked to biological pathways relevant to cave adaptation, we performed Gene Ontology (GO) term analysis. GO term analysis of the 287 BAA genes revealed presence of genes that contribute to metabolic pathways like “chondroitin sulfate proteoglycan metabolic process”, “pyrimidine nucleoside monophosphate metabolic process”, and “pyrimidine nucleoside monophosphate biosynthetic process”. The presence of genes that contribute to pyrimidine-related pathways is consistent with prior findings that intermediates like orotic acid respond sharply to starvation in *A. mexicanus* liver tissue ^4^. Additionally, the BAA genes enriched for non-metabolic terms as well, suggesting a broader systemic change reflected in the genomic compartmental switches of the blind cavefish, extending beyond the liver and metabolism.

### Morph-specific contact loops in surface and cave morphs of *Astyanax*

While A/B compartment analysis provides a broad view of active and inactive genomic regions, understanding gene regulation requires a higher-resolution approach. At a finer scale, we examined chromatin looping, which facilitates enhancer-promoter interactions critical for gene regulation. Using the SIP caller ^36^, we identified 2,525 loops in surface fish, 3,459 in Pachón, and 1,690 in Tinaja, with ∼1,500 loops shared across all morphs. However, 232 loops were unique to surface fish, and ∼200 were common to both cave morphs but absent in surface fish, thereby defining them as ‘cave-specific’. Bullseye metaplots confirmed strong morph-specific signal enrichment at these loops, suggesting regulatory interactions unique to each environment (Fig. 4a). Interestingly, despite these differences in loop composition, the orientations of putative CTCF motifs at the loop anchors remained predominantly convergent across all morphs, suggesting that the mechanisms stabilizing these loops are largely conserved, regardless of morph-specific regulatory changes (Fig. 4b).

**Figure 4:**
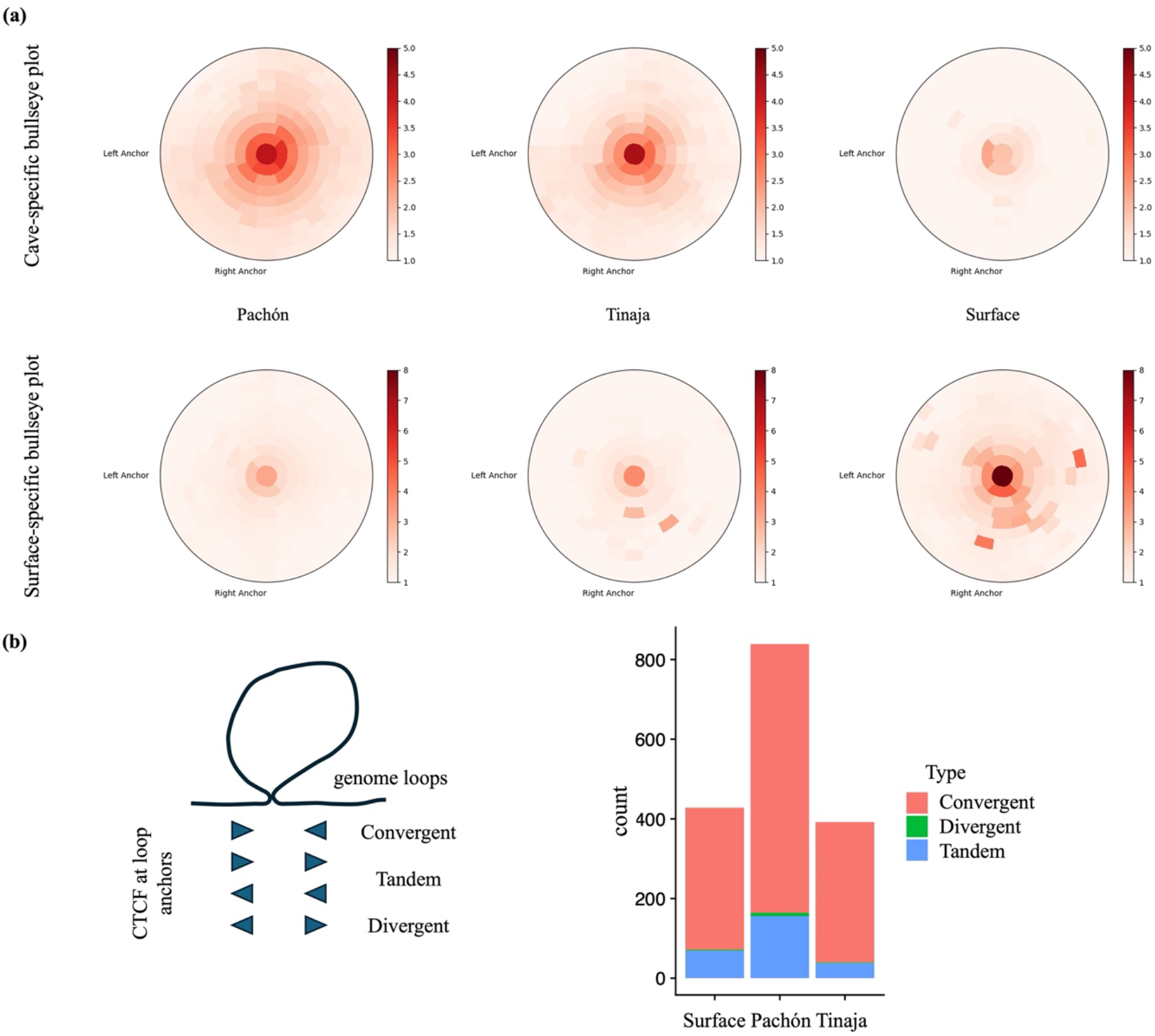
(a) Bullseye transformed visualization from SIPMeta plots for cave-(top) and surface-specific (bottom) loops. (b) Distribution of CTCF anchor points for loops in each of the three morphs. The CTCF sites were either convergent (Surface: 330; Pachón: 677; Tinaja: 456 loops), in tandem (Surface: 55; Pachón: 167; Tinaja: 47 loops) or divergent (Surface: 2; Pachón: 12; Tinaja: 3 loops), as depicted in the cartoon.

### Altered transcriptomic profile under morph-specific loop

Next, we integrated the Hi-C data with RNA-seq, ATAC-seq, and histone ChIP-seq datasets to assess the transcriptional impact of these loops. Of the 620 genes in cave-specific loops, 117 were differentially expressed (p ≤ 0.01), with 75 upregulated in cavefish (Supp. Table 2), while 146 of 667 genes in surface-specific loops were differentially expressed, with 102 upregulated in surface fish (Supp. Table 2).

Loop anchor points, which are often enriched with genes, enhancers and promoters, function as key regulatory regions ^37, 38^. Among the genes upregulated in cavefish within cave-specific loop intervals, we identified two notable candidates: *Arhgef19* (Rho guanine nucleotide exchange factor 19) (Fig. 5a) and *Endog* (endonuclease G) (Supplementary Fig. 3), both positioned near anchor points of cave-specific loops. Notably, *Arhgef19*, which exhibited the highest fold change among these genes, was located near a cave-specific loop anchor. This loop connected to an active enhancer located near the opposite anchor point, as evidenced by H3K27ac and ATAC-seq peaks, suggesting a cave-specific regulatory mechanism (Chr 13:36,235,001-36,275,000) (Fig. 5b). The gene *Endog* was located near the anchor point of a much larger cave-specific loop, approximately 145kb in length (12:1,555,001-1,700,000), on chromosome 12 (Supplementary Fig. 3). Interestingly, a recent collaborative study from our group found that inhibiting the fatty acid transporter *slc27a2a* rescued liver steatosis and atrophy in zebrafish embryos starved for 48 hours ^5^. Interestingly, *Arhgef19* and *Endog* were among the genes differentially expressed between zebrafish subjected solely to starvation and those starved but subsequently rescued. Additionally, both genes have been implicated in adipocyte differentiation and metabolic response to high-fat diets ^39-41^.

**Figure 5:**
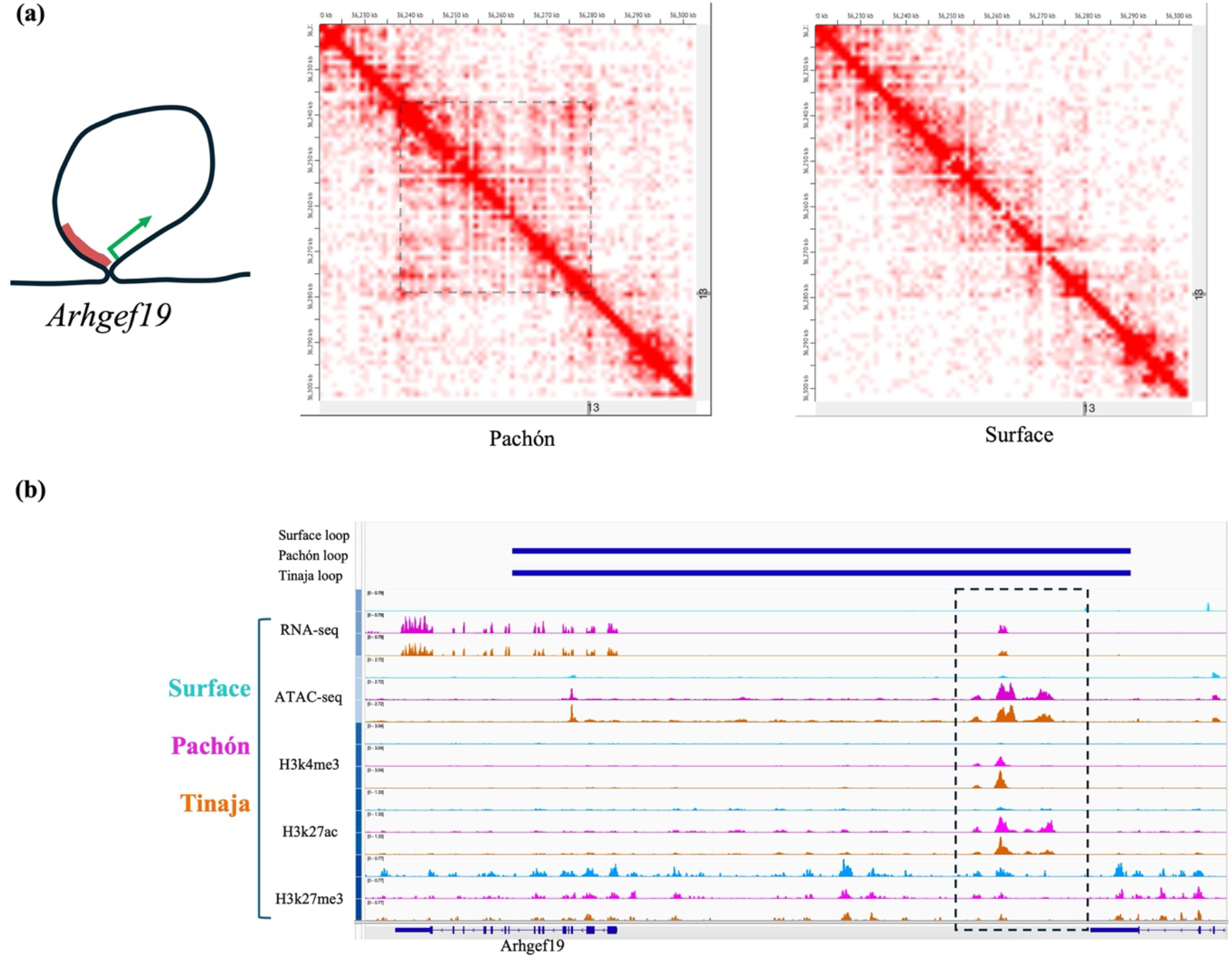
(a) Cave-specific loop of *Arhgef19*, here depicted in Pachón and surface in chromosome 13. Dashed box marks the loop in the Pachón contact matrix. (b) Integrated aligment of the RNA-seq, ATAC-seq and signature histone ChIP-seq data with the loop calls (thick blue lines at the top track depict the span of the loops), to visualize transcriptional activity and regulatory landscape around the loop anchor points. The box marks the putative regulatory region near the other anchor point of the 40kb *Arhgef19* loop.

### Hi-C contact maps reveal inversions in the cavefish genome

Cavefish genomes exhibit structural variations with respect to surface fish and several instances of insertions and deletions were identified across multiple cave morphs, with average cumulative size of cave morph deletions and insertions being 19.4 Mb ^42^. Inversions, which are another major type of structural variation, are known to affect recombination rates and contribute to fitness differences across ecotypes evolving under distinct environmental pressures ^43, 44^. Hi-C’s ability to capture 3D chromatin interactions makes it a powerful dataset for detecting structural variations. Inversions alter expected chromatin contact patterns and disrupt interaction profiles of nearby locations around inversions, often creating a characteristic ‘butterfly’ pattern ^27^ in Hi-C contact maps. To detect inversions in surface and cavefish, we analyzed our Hi-C data using the deep-learning-based EagleC pipeline ^45^, identifying 37 genomic inversions across surface and cave morphs.

One notable inversion, spanning ∼2.7 Mb inversion on chromosome 1 (Chr 1:23,310,000-26,040,000), was shared by both cave morphs (Fig. 6). This inversion had not been previously reported, despite the availability of high-resolution genome assemblies. Notably, the affected region contains several genes involved in cell cycle regulation (Supp. Table 3) – a crucial process for resilience to physiological stress and prevention of tissue atrophy ^5^. Therefore, while a detailed analysis of all inversions is warranted, this inversion itself represents structural variantion through genomic rearrangements as a putative contributor towards resilience to metabolic stresses and cavefish adaptation in general.

**Figure 6:**
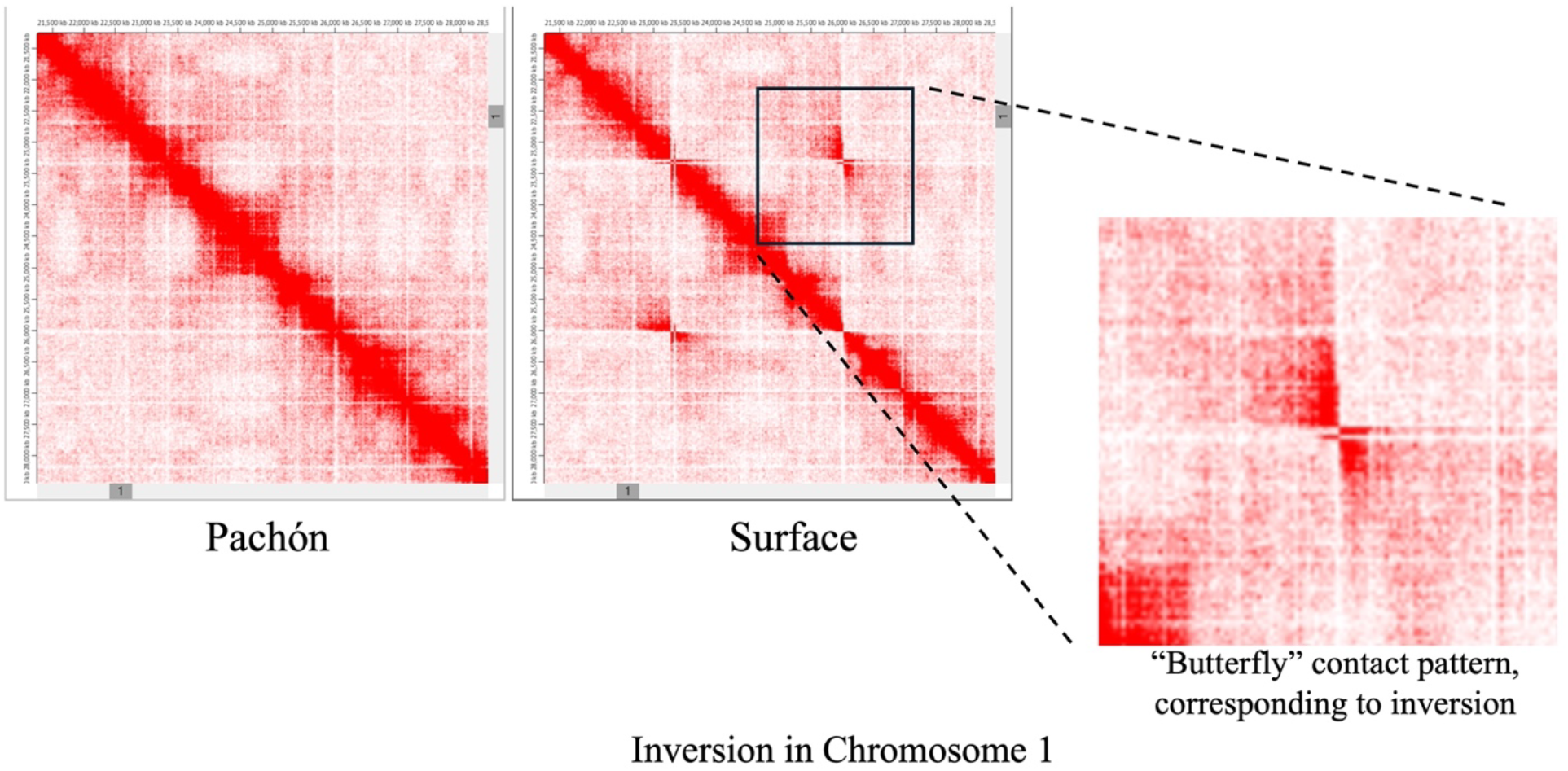
A genomic inversion in between morphs of *Astyanax* in chromosome 1, found in both Pachón and Tinaja with respect to the surface fish (here depicted only comparing Pachón and surface fish). Inset highlights the signature “butterfly” pattern in the Hi-C contact map signifying genomic inversion. To be noted that the inversion appears on surface contact matrix, since the reference is Pachón genome (AMEX_1.1).

## Discussion

In this study, we investigated the 3D genome architecture in liver tissues of surface and cave morphs of *Astyanax mexicanus*, focusing on compartmental organization and chromatin looping to identify structural differences between morphs. Despite belonging to the same species, we uncovered substantial 3D genome differences between surface and cave morphs, with cave-specific features potentially linked to metabolic adaptation, underscoring the role of altered genome architecture in cave-adaptation.

Pathway analysis of genes located within cave-specific 3D compartments and loops revealed a strong enrichment for metabolic processes, suggesting that the 3D genome architecture in *A. mexicanus* livers is primarily tissue-specific. Our Hi-C contact matrix also enabled the identification of long-range regulatory elements, providing a complementary perspective to our previous cis-regulatory study, which focused on proximal regions within 10 kb of target genes ^11^. Through loop analysis, we identified candidate genes such as *Arhgef19* and *Endog*, positioned at cave-specific loop anchors, alongside structural features like inversions, providing novel insights into the genomic basis of cavefish adaptation.

It should also be noted, that while many genes in loci of altered 3D organization showed transcriptomic changes, many genes showed no immediate transcriptional changes (only 75 genes showed higher expression in cavefish, out of 620 genes located in cave-specific loops). A plausible explanation for this is that not all 3D looping variations result in, or are associated with, an immediate effect on gene expression regulation. Instead, these architectural changes may prime regulatory elements for rapid responses to stress, enhancing adaptability ^46^. Further investigation using *in vivo* and *in vitro* ^47, 48^ models for metabolic assays targeting candidate genes within these altered loci will help elucidate their role in adaptability.

The genome looping anchor points, A/B boundaries, and inversion points identified in this study will also become critical for understanding trait evolution, as these are associated with genomic instability and recombination hotspots ^26^, which are known to promote structural variations and influence trait evolution. While QTL (Quantitative Trait Loci) analysis identifies genomic regions linked to adaptive traits, integrating Hi-C data – including loop anchors, inversion points, and A/B compartment boundaries – with the steadily growing list of robust QTL markers will, in the future, help us understand how 3D genome architecture facilitates or constrains evolutionary change, offering insights into the structural basis of cavefish adaptation.

Finally, unlike most 3D genome studies, which compare different species, our work examines morphs within a single species adapted to vastly different environments, laying a foundation for better understanding of how 3D genome changes contribute to speciation and phenotypic diversity. This work not only advances our understanding of *A. mexicanus* evolution but also highlights the broader significance of genome architecture in phenotypic diversity within a single species.

## Acknowledgement

The authors would like to thank Kate Hall and Amanda Lawlor from the Stowers Sequencing and Discovery Genomics team for all the help with Library Preparation and Sequencing. We would also like to acknowledge the University of Kansas Medical Center’s Genomics Core for their support in generating data on the Illumina NovaSeq 6000 System. The core is supported by the following grants: Kansas Intellectual and Developmental Disabilities Research Center (NIH U54 HD 090216), the Molecular Regulation of Cell Development and Differentiation – COBRE (P30 GM122731-03) and the NIH S10 High-End Instrumentation Grant (NIH S10OD021743). This study is supported by Stowers Institute for Medical Research’s institutional funding, and NIH New Innovator Award 1DP2AG071466-01, NIH R24OD030214.

## Author contribution

**Tathagata Biswas**: Conceptualization; data collection; investigation; analysis; visualization; original draft preparation. **Hua Li**: Analysis; original draft preparation, visualization; data curation. **Nicolas Rohner**: Conceptualization; original draft preparation; funding acquisition.

## Data availability statement

The Hi-C data sets has been uploaded to GEO database with accession number GSE290882 (https://www.ncbi.nlm.nih.gov/geo/query/acc.cgi?acc=GSE290882). Original data underlying this manuscript can be accessed from the Stowers Original Data Repository at http://www.stowers.org/research/publications/libpb-2508.

## Supplementary Figures

**Supplementary Fig. 1:**
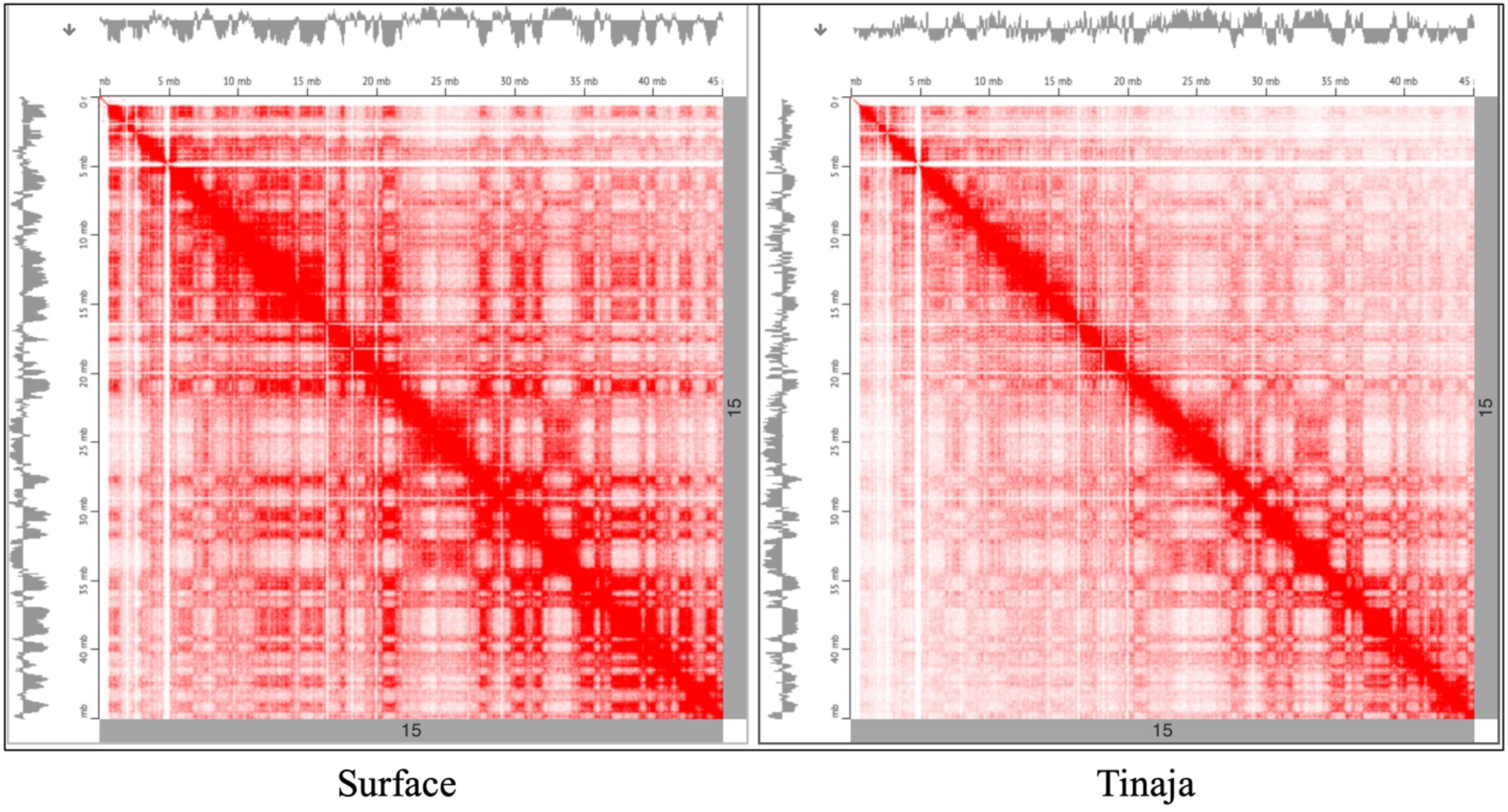
Plaid-like compartment and A/B compartment track for chromosome 15 in Surface and Tinaja.

**Supplementary Fig. 2:**
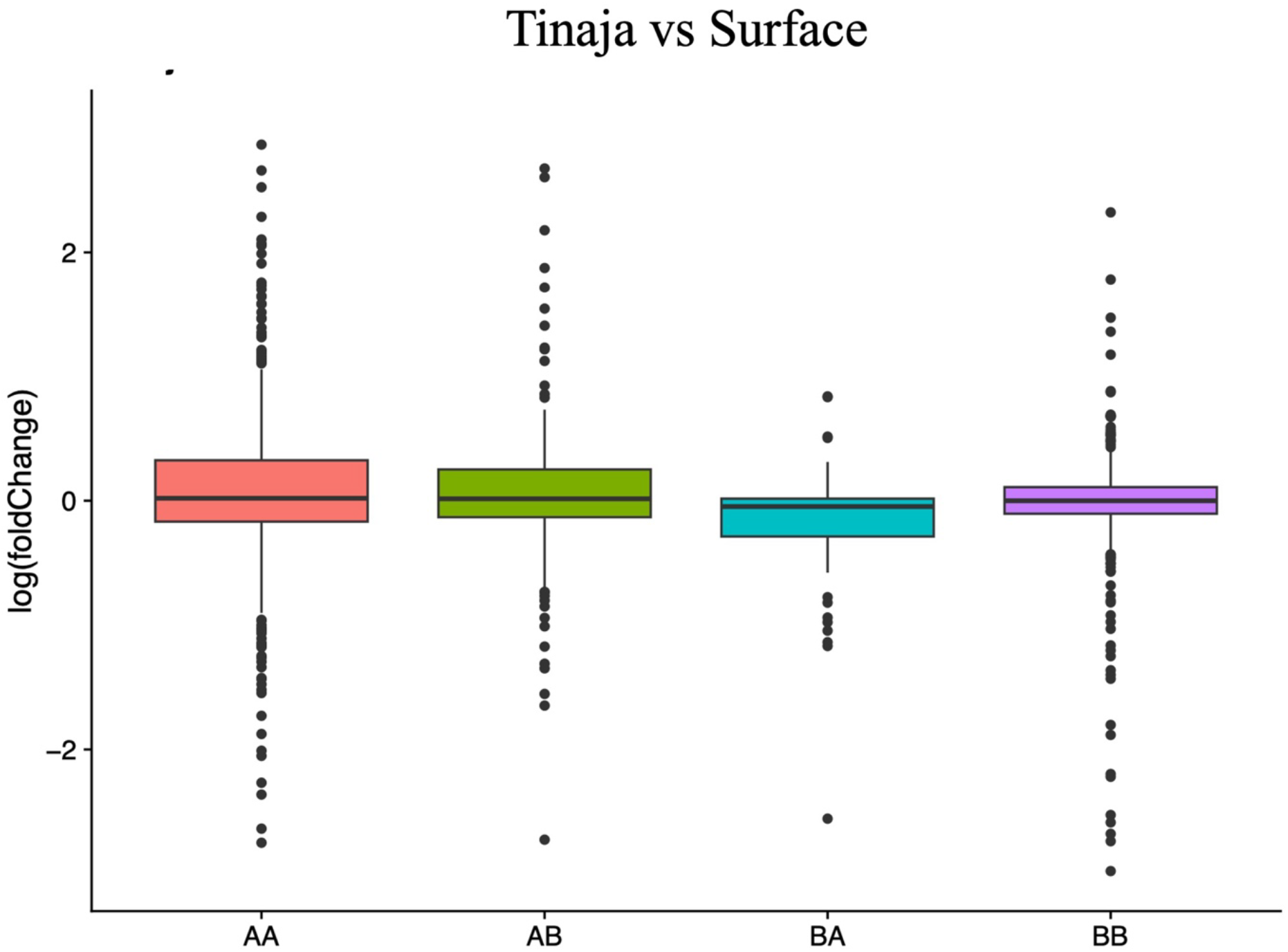
Boxplot depicting fold change in Tinaja, for the genes in the loci which switched compartments. AA and BB denotes unchanged compartments, AB denotes loci which is active in Tinaja and inactive in surface, BA denotes loci active in surface and inactive in Tinaja.

**Supplementary Fig. 3:**
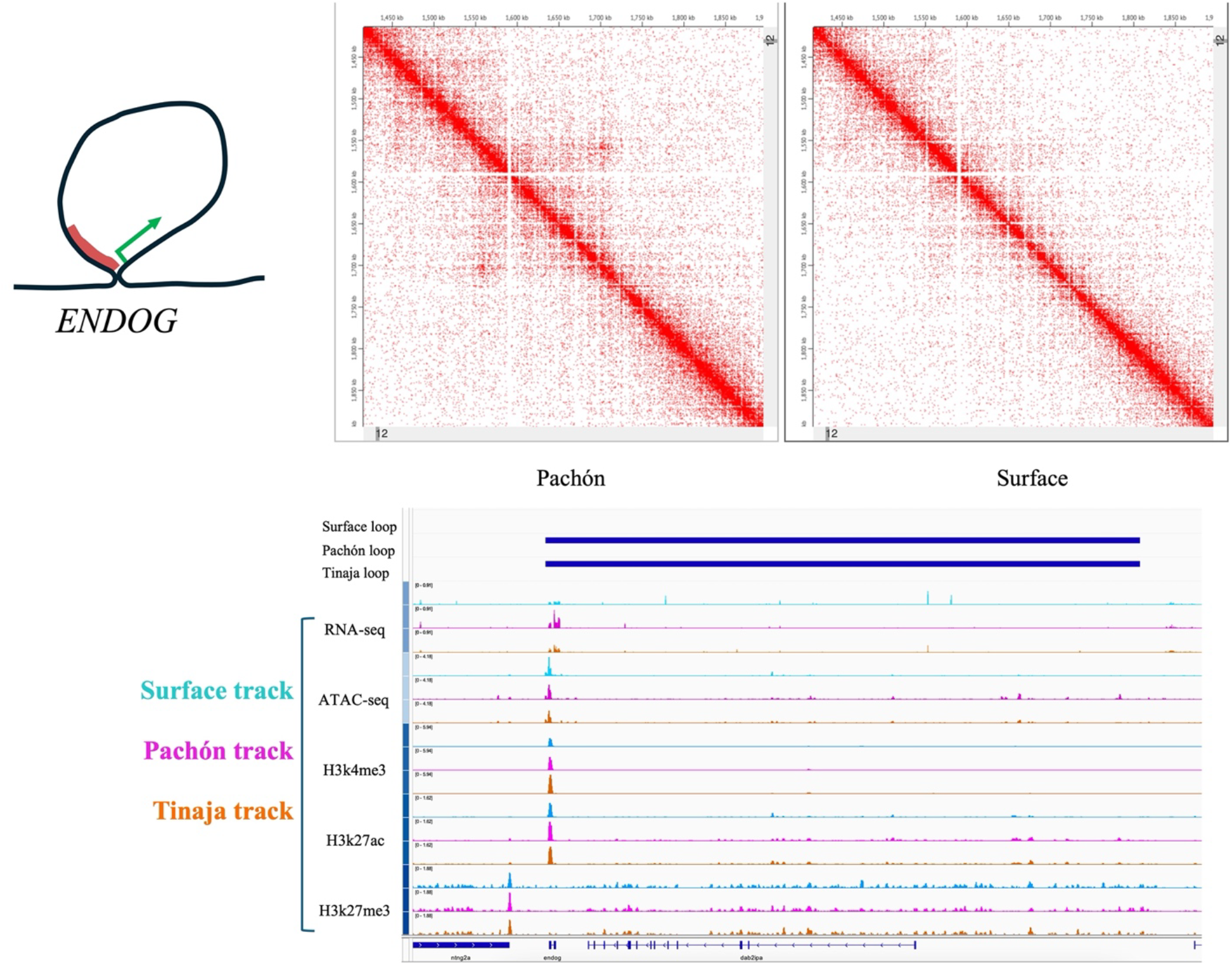
Cave-specific loop, with *Endog* at one of the anchor points. Depicted in Pachón and surface on chromosome 12, integrated with RNA-seq, ATAC-seq and ChIP-seq data.

